# A proteomic signature of vascular dysfunction linked to tauopathy and degeneration in the aging brain

**DOI:** 10.1101/2025.10.27.684881

**Authors:** Hannah L. Radabaugh, Jonah N. Keller, Caleb H. Radtke, Claudia Kunney, Nikolaos Karvelas, Harrison W. Chan, Nivetha Brathaban, Anke Meyer-Franke, Kaitlin Casaletto, Henrik Zetterberg, Bruce L. Miller, Carlos Cruchaga, Joel Kramer, Katerina Akassoglou, Adam R. Ferguson, Fanny M. Elahi

## Abstract

Small vessel disease (SVD) impacts healthy aging of organs across the body, yet its contributions to adverse brain aging remain poorly defined. Here we show thromboinflammation, a core feature of SVD, as a driver of adverse brain aging. We identify cerebrospinal fluid fibrinogen as a marker of brain thromboinflammation and screen neurovascular biosignatures mediating its impact on synaptic vulnerability along the full spectrum of brain aging from cognitively typical, amyloid-negative to cognitively impaired, amyloid-positive older adults. We identified 53 proteins mediating fibrinogen’s effects on synaptic markers in 1,655 donors from three independent cohorts. Single-cell transcriptomic mapping revealed mediator enrichment in neurovascular unit cells. Pathway analysis demonstrated dysregulation of angiogenesis, fibrosis, and immune signaling. Vascular and microglial-enriched biosignatures associated with compromised white matter integrity. These findings indicate thromboinflammation as an early, amyloid-independent pathway to neurodegeneration and tauopathy, establishing vascular health as fundamental to preserving brain healthspan.

## Introduction

Cognitive impairment affects over 57 million people globally, with its prevalence projected to triple by 2050. This growth will create unprecedented public health and socioeconomic challenges, as annual costs rise from $1.3 trillion in 2019 to $2.8 trillion by 2030^1–3^. While age represents the strongest risk factor for all neurodegenerative disorders, current approaches fail to explain this fundamental relationship. Classical neurodegenerative disease biomarkers, including amyloid-β (Aβ) and hyperphosphorylated tau (pTau), capture only 20-40% of variance^4,5^, leaving the majority of age-related cognitive impairment unaccounted for. This critical knowledge gap limits early intervention strategies to preserve brain health and underscores the urgent need to identify drivers of cognitive vulnerability beyond traditional proteinopathy-based models.

Cerebral small vessel disease (cSVD) represents a promising yet understudied contributor to decline in brain function and presents as a common co-pathology across neurodegenerative diseases. Central to cSVD is the development of thromboinflammation in aging blood vessels— a progressive deterioration characterized by increased pro-coagulation states, abnormal endothelial-immune cell interactions, and brain barrier dysfunction^6,7^. This vascular compromise facilitates plasma protein extravasation into the central nervous system (CNS), thereby triggering neuroinflammation through the activation of brain-resident immune cells^8,9^.

Within this cascade, fibrinogen occupies a uniquely central position, acting simultaneously as a coagulation factor and a potent inflammatory signal upon infiltrating the brain parenchyma^10^. Importantly, unlike brain-derived proteins, cerebrospinal fluid (CSF) fibrinogen originates exclusively from peripheral blood, making its quantification a specific indicator of brain barrier dysfunction. Once it infiltrates through compromised barriers, fibrinogen promotes neurovascular unit dysfunction through multiple pathways: neurotoxic microglial programming^11^, oxidative stress induction^12^, and synaptic degeneration^13,14^. Moreover, recent unbiased proteomic analyses provide compelling empirical support for fibrinogen’s role in brain aging^15^. Through an unbiased examination of CSF to plasma ratios across 2,304 peripherally expressed proteins in over 2,000 individuals, fibrinogen exhibited one of the earliest age-related increases in ratio, and was remarkably one of only 10 proteins that were concordantly associated with both aging and cognitive impairment^15^.

Several critical knowledge gaps have limited the translation of insights regarding CNS thromboinflammation into effective interventions for brain aging and cognitive decline. While robust animal studies demonstrate fibrinogen’s neurotoxic effects^6,10–12,14,16,17^, comparable human investigations remain largely absent. Furthermore, the specific molecular networks connecting vascular dysfunction at brain barriers to neuronal damage within brain parenchyma remain uncharacterized. The temporal relationship between thromboinflammation and the accumulation of amyloid pathology in the aging brain is similarly unknown. Addressing these gaps is critical for developing vascular-targeted biomarkers and interventions that could preserve brain health.

We present the largest investigation of thromboinflammation in the aging human brain, analyzing high-dimensional CSF proteomics data (∼7,000 proteins) across 1,655 participants aged 37 to 91 years from three independent cohorts. Our systematic approach provides unprecedented characterization of thromboinflammatory networks in the CSF of living human brains. By using CSF molecular indices of neurodegeneration, we identified specific molecular mediators linking vascular dysfunction to neuronal vulnerability and revealed signatures that operate independently and possibly upstream of amyloid pathology. Our findings illuminate vascular-directed therapeutic targets and could potentially inform early intervention strategies for brain health preservation, addressing a critical unmet need in exponentially aging populations.

## Results

To systematically characterize the molecular pathways driving age-related brain vulnerability to degeneration, we analyzed high-dimensional CSF proteomics data (7,596 proteins via SOMAscan 7k platform) across 1,655 individuals from three independent cohorts, spanning normal cognition to impaired. Of these, 638 individuals were amyloid-negative, cognitively normal older adults aged 46 to 91 years (**Extended Data Fig. 1**; **Supplementary Table 1**). To ensure the relevance of our discoveries to early, pre-amyloid aging, we began our analyses in this subgroup. The deeply-phenotyped discovery cohort (UCSF Hillblom) enabled initial pathway identification, while two additional validation cohorts provided independent replication across geographically diverse populations. In this initial analysis, all participants presented with typical cognitive function for age, no functional impairment (Clinical Dementia Rating Sum of Boxes = 0), and demonstrated absence of amyloid pathology (18F-AV-45 SUVR ≤ 1.32^18^ or PHC Centiloids < 25^19^). Technical validation using proteomics of duplicate CSF aliquots confirmed robust analytical precision (coefficients of variation < 20% for 95% of proteins; **Extended Data Fig. 2a-d**). Once we established a reliable foundation for cross-cohort molecular comparisons, we tested the persistence of findings in amyloid-negative aging (*N* = 638), in amyloid-positive individuals (*N* = 1017; **Fig. 1**).

**Figure 1.**
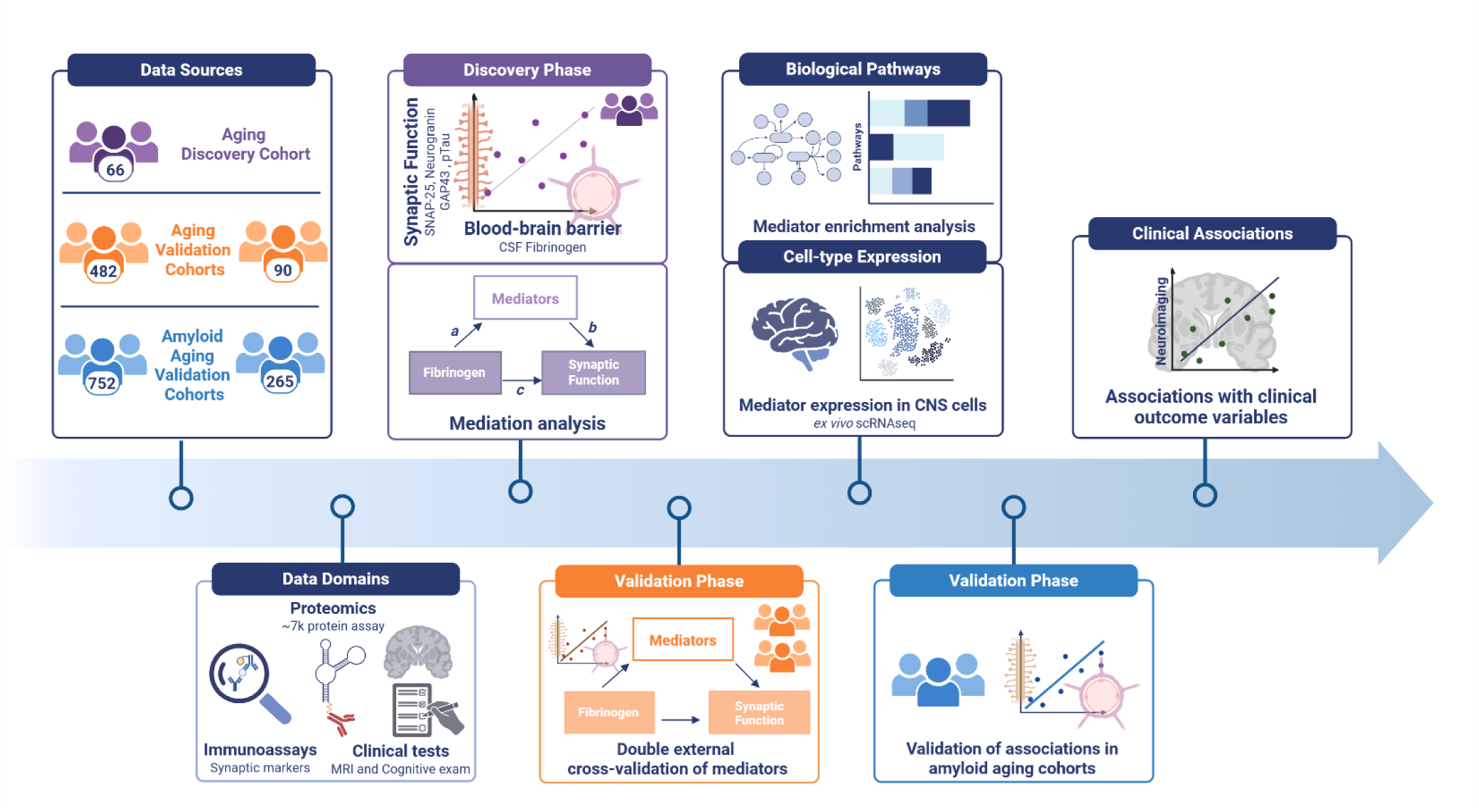
Study Design. A comprehensive, graphical representation of the analytical workflow applied throughout the current study. Icons on all subsequent figures redirect to phases outlined here. Colors correspond to specific cohorts throughout the manuscript with purple referring to the Aging Discovery Cohort, orange referring to Aging Validation Cohorts and blue referring to Amyloid Aging Validation Cohorts.

### Protein co-expression network analysis in aging CSF identifies a conserved thromboinflammatory module centered on fibrinogen

Weighted protein co-expression network analysis (WPCNA) of CSF proteomic data identified a distinct 116-protein co-expression module significantly enriched for thromboinflammatory proteins across all cohorts^20^. After assessing patterns of missingness and imputing missing values (**Fig. 2a,b**), fibrinogen emerged as a hub gene of this module in the discovery cohort. Replication in independent cohorts showed stronger module membership in Knight ADRC and ADNI, with high network preservation scores indicating robust maintenance of co-expression relationships across aging populations (**Fig. 2c**).

**Figure 2.**
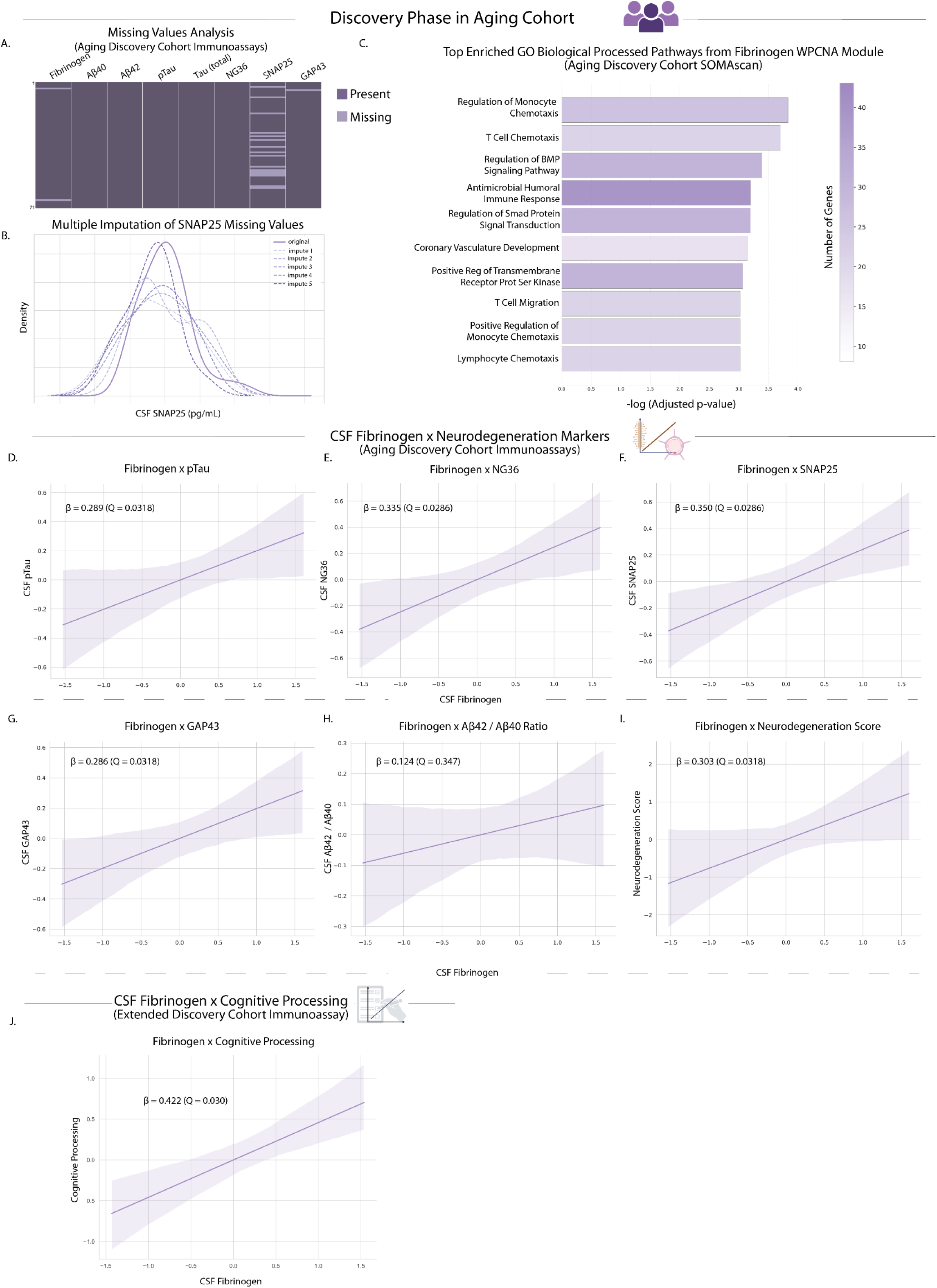
Elevated CSF fibrinogen levels are associated with increased markers of neurodegeneration independent of amyloidopathy. **a,** Patterns of missingness of targeted immunoassay values in the discovery cohort (light purple indicates missing; 2.8% for fibrinogen, 22.5% for SNAP25; both missing completely at random) and **b,** multiple imputation of missing SNAP25 values. **c,** Weighted protein co-expression network visualization of the distinct 116-protein thromboinflammatory module. Module membership scores reflect the correlation between fibrinogen’s expression pattern and the module’s overall expression profile (eigengene): discovery cohort (0.36, *P* = 0.003), Knight ADRC (0.73, *P* = 1.59e-80), ADNI (0.58, *P* = 2.44e-7). Network preservation scores (0.85 and 0.96) indicate robust maintenance of pairwise co-expression relationships within the module across cohorts. Pathway enrichment analysis showed significant overrepresentation of: regulation of monocyte chemotaxis (*P* = 1.47e-4), T cell chemotaxis (*P* = 1.98e-4), lymphocyte chemotaxis (*P* = 9.54e-4), and antimicrobial humoral immune responses (*P* = 6.33e-4). **d-g,** Regression plots showing associations between CSF levels of fibrinogen and markers of synaptic degeneration in cognitively typical, amyloid-negative adults (*N* = 66): pTau181 (β = 0.289, *Q* = 0.032), NG36 (β = 0.335, *Q* = 0.029), SNAP25 (β = 0.350, *Q* = 0.029), and GAP43 (β = 0.286, *Q* = 0.032). **h,** No significant association was observed between fibrinogen and Aβ42/40 ratio (β = 0.124, *Q* = 0.347). **i,** A principal component analysis of synaptic biomarkers yielded a Neurodegeneration Score (PC1, 86.5% variance explained), which was significantly associated with CSF fibrinogen levels (β = 0.303, *Q* = 0.032). **j,** Regression plot showing the association of CSF Fibrinogen in the Extended Discovery Cohort (all subjects enrolled in the UCSF Hillblom cohort regardless of amyloid status) with cognitive processing (ms) (β = 0.422, Q = 0.030). Higher processing time (ms) indicates worse performance on this task. All CSF immunoassay values are log_2_ transformed and all associations are controlled for age and sex.

The module encompassed interconnected pathways of pathophysiological significance, including biglycan and decorin-mediated inflammatory cascades^21,22^, thrombospondin activation^23–25^, PGLYRP1-mediated microglial activation^26,27^, and CXCL chemokine signaling networks^28^. To functionally characterize this module, pathway enrichment analysis revealed significant overrepresentation of immune cell recruitment processes, including regulation of monocyte chemotaxis, T cell chemotaxis, lymphocyte chemotaxis, and antimicrobial humoral immune responses (**Fig. 2c**). These findings demonstrate detectability of thromboinflammatory networks from CSF of typically aging adults, hinting at coordinated activation of vascular and immune pathways during typical brain aging. The central role of fibrinogen in this thromboinflammatory network aligns with recent findings which identified fibrinogen as a peripheral protein exhibiting one of the earliest age-related increases in CSF to plasma ratio^15^.

### Elevated CSF levels of blood fibrinogen positively associate with markers of synaptic dysfunction and neurodegeneration in typically aging adults

Having established fibrinogen as a central hub of thromboinflammatory networks detected in aging CSF, we next tested whether CSF fibrinogen levels associate with markers of synaptic vulnerability in cognitively typical aging. To validate fibrinogen as a specific indicator of brain barrier dysfunction rather than endogenous brain inflammation, we first confirmed its exclusively peripheral origin using single-cell RNA sequencing data from *ex vivo* human brain tissue^29^. Analysis demonstrated that none of fibrinogen’s constituent subunit transcripts (*FGA*, *FGB*, *FGG*) were detected in any brain cell type (**Extended Data Fig. 3a-d**), confirming that CSF fibrinogen specifically reflects brain barrier dysfunction and extravasation of blood components into the CNS, consistent with previous reports^6,10,12,14,15,17,30,31^.

Linear regression analyses revealed that higher CSF fibrinogen levels were significantly associated with elevated markers of synaptic degeneration and tauopathy in amyloid-negative older adults, independent of age and sex (**Fig. 2d-i**; **Supplementary Table 2**). Specifically, fibrinogen levels correlated positively with phosphorylated tau (pTau181), neurogranin (NG36), synaptosomal-associated protein 25 (SNAP25), and growth-associated protein 43 (GAP43) **(Fig. 2d-g**). Critically, no significant association was observed between fibrinogen and the Aβ_42/40_ ratio, demonstrating independence from amyloid pathology (**Fig. 2h**). To account for the correlated nature of these synaptic biomarkers, we performed principal component analysis (PCA) to generate a composite Neurodegeneration Score, which showed significant positive association with CSF fibrinogen levels (**Fig. 2i**). These findings demonstrate that thromboinflammation, as quantified by CSF fibrinogen, associates with synaptic vulnerability. The absence of correlation with direct biomarkers of amyloidopathy suggest that these precede amyloid pathology or are independent of it.

Given the association of fibrinogen with biomarkers of neurodegeneration, we asked whether fibrinogen levels also correlate with variance in cognitive processing speed, a cognitive function that declines with age. Testing this hypothesis in the extended discovery cohort, which included all participants with available cognitive processing speed, we found that higher levels of CSF fibrinogen were significantly associated with slower cognitive processing speed (**Fig. 2j**).

### Seventy-two protein mediators link fibrinogen to markers of synaptic injury and neurodegeneration

To uncover the molecular pathways through which thromboinflammation contributes to synaptic injury during typical aging, we conducted comprehensive mediation analyses to identify proteins that mediate the relationship between CSF fibrinogen and synaptic dysfunction markers. Using a rigorous application of the Baron and Kenny framework^32^, controlling for age, sex, and false discovery rate, we screened SOMAmers for their potential mediating role between fibrinogen, as a measure of thromboinflammation, and each synaptic marker (pTau, GAP43, NRGN/NG36, and SNAP25). Statistical stability was ensured through nonparametric bootstrapping with 1,000 iterations per analysis.

This systematic approach identified 74 SOMAmers (72 unique proteins with 3 different SOMAmers each capturing FN1) that consistently mediated the fibrinogen-synaptic biomarker relationships across all four individual screens, representing shared molecular pathways linking thromboinflammation to neurodegeneration in typically aging adults (**Fig. 3a-c**). To account for the intercorrelated nature of synaptic biomarkers, we orthogonally validated these findings using the composite Neurodegeneration Score (PC1). All 74 previously identified SOMAmers maintained statistical significance when tested against this composite measure, further solidifying their central role in thromboinflammation-associated synaptic vulnerability. These validated mediators span diverse biological functions relevant to vascular integrity, immune activation, and synaptic maintenance, providing molecular insights into how systemic subclinical age-associated thromboinflammation may influence brain health (**Fig. 3d**; **Supplementary Table 3**).

**Figure 3.**
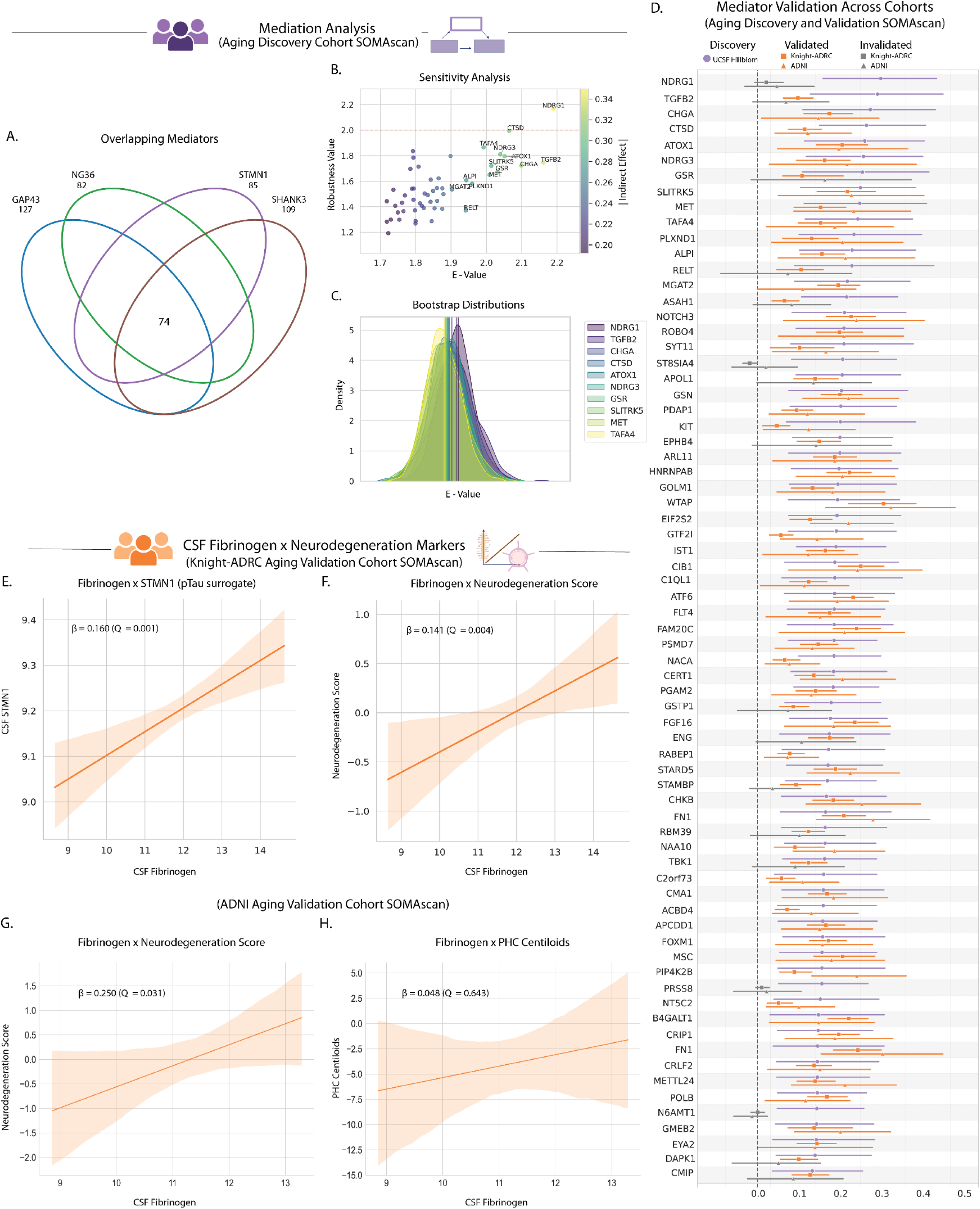
Identification of 53 common protein mediators linking fibrinogen to markers of neurodegeneration. **a,** 74 shared SOMAmers significantly mediate the association between CSF levels of fibrinogen and markers of synaptic degeneration (pTau, GAP43, NRGN, SNAP25) in cognitively typical, amyloid-negative older adults (*N* = 66). **b,** Sensitivity analysis determining robustness of identified proteins. **c,** Mediation effects were estimated using the Baron and Kenny framework with 1,000 bootstrap iterations per test. Cross-platform concordance showed strong agreement between SOMAscan and targeted immunoassay measurements: fibrinogen, GAP43, and NRGN (*Q* < 5.00e-11). Biologically validated surrogate markers: STMN1 for pTau (*Q* = 1.95e-16) and SHANK3 for SNAP25 (*Q* = 5.89e-15). **d,** Of the 74 SOMAmers, 54 (53 unique proteins) replicated in both independent aging cohorts (Knight-ADRC *N* = 482; ADNI *N* = 90). **e,** Representative replication showing fibrinogen association with pTau surrogate STMN1 in Knight-ADRC (β = 0.160, *Q* = 1.49e-3). **f,** Representative replication showing fibrinogen association with Neurodegeneration Score in Knight-ADRC (β = 0.141, *Q* = 3.99e-3). Additional Knight-ADRC replication results (not shown): SHANK3 (SNAP25 surrogate; β = 0.173, *Q* = 1.14e-3), GAP43 (β = 0.112, *Q* = 0.019), and NRGN (β = 0.140, *Q* = 3.99e-3). **g,** Fibrinogen– Neurodegeneration Score association in ADNI (β = 0.250, *Q* = 0.031). Additional ADNI replication results (not shown): STMN1 (pTau surrogate; β = 0.281, *Q* = 0.023), SHANK3 (SNAP25 surrogate; β = 0.229, *Q* = 0.038), GAP43 (β = 0.246, *Q* = 0.031), and NRGN (β = 0.283, *Q* = 0.023). **h,** No association was observed between fibrinogen and cortical amyloid load (PHC Centiloids) in ADNI (β = 0.048, *Q* = 0.643). All CSF levels of proteins are log_2_ transformed and all associations are controlled for age and sex.

### Cross-platform validation of protein signature demonstrates thromboinflammation-synaptic association replication across independent cohorts

To establish the generalizability of thromboinflammation-synaptic relationships, we performed external validation across two independent cohorts of typically aging, amyloid-negative older adults: Knight-ADRC (*N* = 482) and ADNI (*N* = 90). We harmonized datasets by establishing cross-platform concordance between SOMAscan and targeted immunoassay measurements (**Extended Data Fig. 4a-c**; **Supplementary Table 4**). We identified biologically-relevant surrogate markers for pTau (STMN1, a microtubule-associated phosphoprotein^33,34^) and SNAP25 (SHANK3, a synaptic scaffolding protein^35^) (**Extended Data Fig. 4d-g**; **Supplementary Table 4**).

Using these validated cross-platform mappings, we tested whether the discovery-phase thromboinflammation-neurodegeneration relationships replicated in both validation cohorts. Higher CSF fibrinogen levels were robustly associated with elevated pTau (measured through the surrogate STMN1) and synaptic dysfunction markers in both Knight-ADRC and ADNI cohorts (**Fig. 3e**; **Extended Data Fig. 5a-h**; **Supplementary Table 4**). The composite Neurodegeneration Score showed consistent associations across cohorts (**Fig. 3f,g**; **Supplementary Table 4**). Critically, fibrinogen again showed no association with amyloid burden (PHC Centiloids) in ADNI, confirming amyloid independence (**Fig. 3h**; **Supplementary Table 4**). These findings establish the reproducibility of thromboinflammation-synaptic vulnerability relationships across diverse populations and measurement platforms.

### Fifty-three of seventy-two protein mediators of thromboinflammation were externally validated in two independent aging cohorts

Mediation screening in validation cohorts using fibrinogen (SOMAmer SeqID: 4907-56) as input and the composite Neurodegeneration Score (derived from SHANK3 [SeqID: 13242-134], GAP43 [SeqID: 15325-14], STMN1 [SeqID: 17367-5], and NG36/NRGN [SeqID: 18303-39]) as output identified 54 SOMAmers representing 53 unique proteins that consistently mediated thromboinflammation-synaptic vulnerability relationships. These validated mediators demonstrated robust statistical significance across all cohorts, discovery (UCSF) and validation (ADNI and Knight-ADRC), establishing a reproducible molecular network linking vascular barrier compromise to synaptic vulnerability in typically aging adults (**Fig. 3d**; **Supplementary Table 3,4**). The high replication rate (54 of 74 SOMAmers, 73%) across independent populations suggests generalizability of this mediator network. The 20 SOMAmers that failed validation criteria were excluded from subsequent analyses to ensure only the most robust findings informed downstream functional characterization, further limiting the risk of false discovery inherent to high-dimensional data analyses.

### Validated protein mediators encompass three coordinated biological networks linking thromboinflammation to synaptic vulnerability

The 53 validated protein mediators revealed distinct functional networks that mechanistically connect vascular barrier compromise to synaptic dysfunction. A prominent vascular integrity and endothelial function cluster included several proteins with established roles in cerebral small vessel disease. *NOTCH3*, encoding a mural-cell receptor, is mutated in CADASIL, a monogenic form of cSVD leading to early-onset stroke and dementia. *FN1* emerged as a particularly intriguing mediator, as it is upregulated in both sporadic and genetic forms of cSVD^36–38^, yet rare loss-of-function variants in *FN1* protect against Alzheimer’s disease in *APOE4* homozygotes^38,39^. This relationship suggests that *FN1* may represent a critical mediator linking vascular pathology to neurodegeneration. The cluster also included *FLT4* which regulates angiogenic functionality and has documented involvement in both Alzheimer’s disease^40^ and genetic cSVD^37^, as well as *PLXND1*, a mechanosensor that enables endothelial cells to detect shear stress and regulate vascular responses^41^.

Another prominent inflammatory and fibrotic signaling cluster included *CTSD*, a lysosomal protease that is upregulated in plaque-associated microglia and represents a key marker of microglial activation in Alzheimer’s disease^42^. Other proteins in this network, such as *ATF6* and *B4GALT1*, regulate ceramide balance, endoplasmic reticulum stress, and integrin glycosylation, respectively^43,44^.

The third cluster comprised synaptic and neuronal signaling proteins, including complement component *C1QL1*, which couples oligodendrocyte maturation to white matter integrity^45,46^, *SLITRK5*, a brain-derived neurotrophic factor co-receptor organizing corticostriatal synapses^47^, and *SYT11* which regulates vesicle cycling in dopaminergic terminals^48,49^.

This coordinated network architecture reveals how thromboinflammation might propagate vascular dysfunction through neuroinflammatory cascades that ultimately compromise synaptic integrity.

### CSF enrichment analysis reveals pathways indicative of brain barrier compromise alongside activation of vascular, immune, and proliferative processes

To gain deeper insights into the biological pathways represented by the 53 validated mediator proteins, we performed comprehensive pathway enrichment analyses using EnrichR across eight curated libraries (**Fig. 4a,b**; **Supplementary Table 5**). When analyzing mediators by their directional associations with neurodegeneration, distinct pathway signatures emerged.

**Figure 4.**
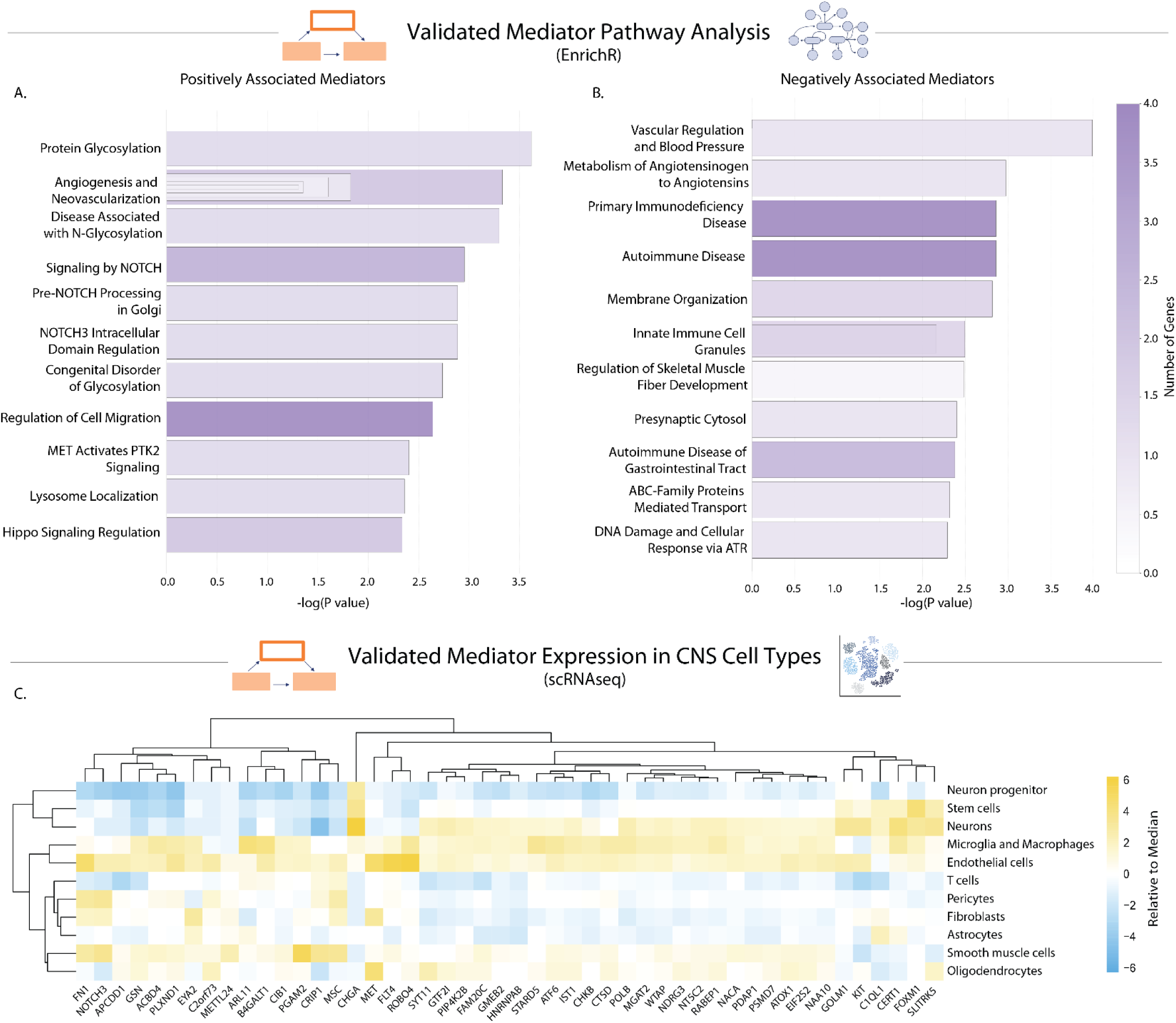
Pathway and single cell expression analysis of 53 validated thromboinflammatory mediators reveals central role of brain barrier compromise and neurovascular unit. Pathway analysis of the validated thromboinflammatory mediators using the EnrichR platform incorporating eight curated libraries: GO Biological Process (2025), GO Cellular Component (2025), GO Molecular Function (2025), Jensen (Disease Curated and Experimental; 2025), Reactome (2024), SynGO (2024), and WikiPathways (2024). **a,** Mediators positively associated with neurodegeneration (31 genes) showed enrichment for neovascularization processes (*P* = 4.26e-4), protein glycosylation (*P* = 2.27e-4), and NOTCH signaling pathways (*P* = 1.22e-3). **b,** Mediators negatively associated with neurodegeneration (21 genes) were enriched for vascular regulation (*P* = 1.03e-4), synaptic processes (*P* = 3.97e-3), ficolin-1-rich granule components (*P* = 3.20e-3 for lumen, *P* = 6.90e-3 for granules), and autoimmune disease pathways (*P* = 1.38e-3). **c,** One-versus-rest differential-expression analysis of an *ex vivo* scRNA-seq atlas across principal neurovascular and glial cell types. Endothelial cell-selective expression: *FLT4* (log₂FC = 7.29, 15.8% of cells), *ROBO4* (5.54, 72.2%), *PLXND1* (2.07, 44.0%). Multi-cellular patterns: *FN1* in pericytes (3.53, 86.5%), endothelial cells (3.29, 90.2%), fibroblasts (1.86, 68.6%), smooth-muscle cells (0.85, 64.1%); *GSN* in fibroblasts (1.37, 80.5%), oligodendrocytes (1.29, 93.5%), smooth-muscle cells (0.76, 90.9%), endothelial cells (0.70, 85.2%), microglia/macrophages (0.61, 64.7%), pericytes (0.50, 70.7%). Oligodendrocyte enrichment: *SYT11* (3.12, 95.1%), *ATOX1* (1.72, 76.4%), *NDRG3* (0.98, 23.4%). Mural cell expression: *NOTCH3* in pericytes (4.69, 91.3%) and smooth-muscle cells (2.73, 87.6%); *CRIP1* in pericytes (1.84, 14.0%) and smooth-muscle cells (4.77, 51.7%). Microglial/macrophage enrichment: *B4GALT1* (2.88, 41.3%), *CTSD* (1.32, 42.4%), *ATF6* (1.48, 26.6%).

Positively associated mediators showed the most robust enrichment for neovascularization processes, indicating active microvascular remodeling and angiogenic activity, along with protein glycosylation and NOTCH signaling pathways (**Fig. 4a**; **Supplementary Table 5**). Conversely, negatively associated mediators were enriched for vascular regulation, as well as for synaptic and immune-related processes, including innate immune cell granules and autoimmune disease pathways (**Fig. 4b**; **Supplementary Table 5**). The pathway signatures suggest coordinated vascular remodeling responses that accompany changes in immune function and synaptic vulnerability.

### Protein mediators validated in external cohorts show cell-type specificity within the neurovascular unit

To better understand the cellular sources of the protein mediators of thromboinflammation, we performed one-versus-rest differential-expression analysis using an *ex vivo* scRNA-seq atlas^29^, considering only the validated genes across the principal neurovascular and glial cell types (**Fig. 4c**; **Supplementary Table 6**). Among the vascular constituents, *FLT4* exhibited the most selective expression in endothelial cells, alongside *ROBO4* and *PLXND1*. Several mediators displayed multi-cellular patterns: *FN1* was highly expressed in pericytes, endothelial cells, fibroblasts, and smooth muscle cells, while *GSN* was abundant in fibroblasts, oligodendrocytes, smooth-muscle cells, endothelial cells, microglia/macrophages, and pericytes. Within oligodendrocytes, *SYT11* showed the highest enrichment, followed by *ATOX1* and *NDRG3*.

Among mural cells, *NOTCH3* and *CRIP1* were highly expressed in pericytes, mirroring the smooth muscle signature where they remain prominent. In brain-resident immune cells, microglia and macrophages were enriched for *B4GALT1*, *CTSD*, and *ATF6*. Collectively, these expression profiles indicate that proteins within the thromboinflammatory network are concentrated in endothelial, mural, and immune cell subpopulations, with *FN1* and *GSN* bridging multiple vascular lineages, suggesting a coordinated vascular-immune network that may ultimately modulate synaptic integrity.

### CSF signature of thromboinflammation is relevant to clinical outcomes

We then sought to evaluate whether shared variance among proteins in the thromboinflammatory signature relates to brain structural integrity, known to decline with aging. To define a summary composite score of the mediator profile, we performed PCA on the 54 validated SOMAmers. The first principle component (Mediator PC1) captured the majority of variance (67.9%) across the mediator network (**Fig. 5a**; **Supplementary Table 7**). Moreover, the PC was positively associated with pTau (**Fig. 5b**; **Supplementary Table 8**) and the Neurodegeneration Score (**Extended Data Fig. 6a**; **Supplementary Table 8**). No significant associations were found with amyloid (**Fig. 5c**; **Supplementary Table 8**).

**Figure 5.**
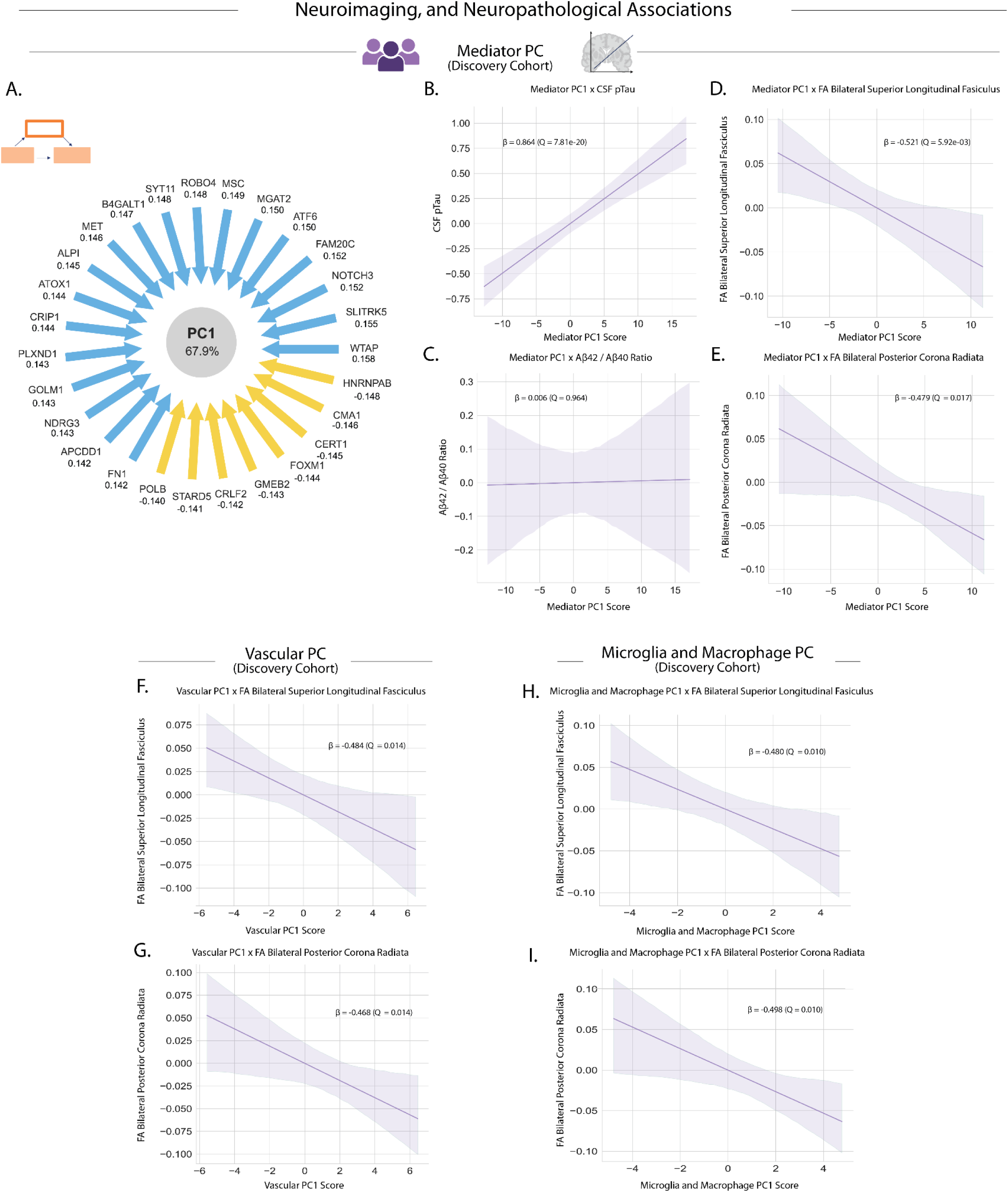
Thromboinflammatory proteins are significantly associated with cognitive, neuroimaging, and neuropathological outcomes. **a,** Mediator PC1 (67.9% variance explained), a composite value of individual mediator protein expression in the discovery cohort, positively associated with Neurodegeneration Score (β = 0.844, *Q* = 6.96e-18). Loading values and relative directionality shown. **b,** Mediator PC1 association with pTau surrogate STMN1 (β = 0.864, *Q* = 7.81e-20). **c,** Mediator PC1 shows no association with Aβ42/40 ratio (β = 0.006, *Q* = 0.964). **d-e,** Associations of Mediator PC1 scores with white matter structural integrity: bilateral superior longitudinal fasciculus fractional anisotropy (FA) values (β = −0.521, *Q* = 5.92e-3) and bilateral posterior corona radiata (β = −0.479, *Q* = 0.017). **f-g,** Vascular PC1 (66.1% variance explained), composite describing proteins enriched in vascular cell types in the discovery cohort, associations with white matter structural integrity: bilateral superior longitudinal fasciculus (β = −0.484, *Q* = 0.014) and bilateral posterior corona radiata (β = −0.468, *Q* = 0.014). **h-i,** Microglia and Macrophage PC1 (74.1% variance explained), composite describing proteins enriched in microglia and macrophages in the discovery cohort, associations with white matter structural integrity: bilateral superior longitudinal fasciculus (β = −0.480, *Q* = 0.010) and posterior corona radiata (β = −0.498, *Q* = 0.010). All associations are controlled for age and sex.

Regression models demonstrated that higher Mediator PC1 scores were significantly associated with compromised microstructural integrity across multiple large white matter tracts, specifically in the bilateral superior longitudinal fasciculus and bilateral posterior corona radiata (**Fig. 5d,e**; **Supplementary Table 8**). These findings suggest that the composite pattern of mediator proteins is associated with reduced microstructural integrity in large white matter tracts with implications for cognitive aging^50,51^.

### Cell-type specific molecular profiles reveal distinct contributions to white matter integrity

To further investigate whether mediators enriched in specific cell-types were relevant to white matter associations, we performed targeted analyses on mediator subsets filtered by cell type, as determined by our single-cell RNA sequencing analysis. Vascular cell-enriched mediators (combining endothelial cells, pericytes, and smooth muscle cells) showed particularly robust associations with white matter integrity in the bilateral superior longitudinal fasciculus and bilateral posterior corona radiata (**Fig. 5f,g**; **Supplementary Table 8**).

Complementary analyses of microglia and macrophage-enriched mediators also revealed significant associations with white matter integrity in the same tracts (**Fig. 5h,i**; **Supplementary Table 8**). These findings underscore the coordinated involvement of multiple neurovascular unit cell types in mediating thromboinflammation-associated brain pathology, with both vascular and immune cell signatures showing strong relationships to white matter damage as an early clinical outcome of the aging brain.

### Subclinical thromboinflammation signatures associate with tauopathy and associations persist into amyloid-positive neurodegenerative disease states

We demonstrated that fibrinogen is associated with pTau in the discovery cohort (**Fig. 2d**) and replicated these associations in both Knight-ADRC and ADNI typical aging cohorts (**Fig. 3e**; **Extended Data Fig. 5a,e**; **Supplementary Table 4**). Critically, fibrinogen showed no association with amyloid burden in typical aging in either the discovery cohort (**Fig. 2h**) or ADNI validation cohort (**Fig. 3h**), confirming independence from amyloidopathy. Similarly, the mediator PC1 composite score demonstrated strong associations with pTau while maintaining amyloid independence (**Fig. 5b,c**; **Supplementary Table 8**).

To further investigate this amyloid independence, we assessed the relevance of thromboinflammation to amyloid-positive disease states by conducting comprehensive analyses in amyloid-positive participants from both validation cohorts. This investigation was designed to test the hypothesis that fibrinogen-associated thromboinflammatory pathways act through mechanisms fundamentally distinct from the classical amyloid cascade hypothesis, spanning the continuum from typical brain aging to proteinopathy-associated neurodegeneration.

Linear regression analyses in amyloid-positive cohorts revealed robust preservation of thromboinflammation-neurodegeneration associations. In the Knight-ADRC amyloid-positive cohort (*N* = 752), fibrinogen demonstrated significant associations with the pTau surrogate STMN1 (**Fig. 6a**; **Supplementary Table 9**) and the composite Neurodegeneration Score (**Fig. 6b**; **Supplementary Table 9**), with effect sizes comparable to amyloid-negative populations (**Extended Data Fig. 7a-c**; **Supplementary Table 9**). Independent validation in the ADNI amyloid-positive cohort (*N* = 265) confirmed these findings with maintained statistical significance and comparable effect sizes (**Fig. 6c,d**; **Extended Data Fig. 7d-f**; **Supplementary Table 9**). Moreover, while fibrinogen maintained its association with tauopathy in amyloid-positive ADNI participants, there was no significant relationship with amyloid (**Fig. 6e**; **Supplementary Table 9**).

**Figure 6.**
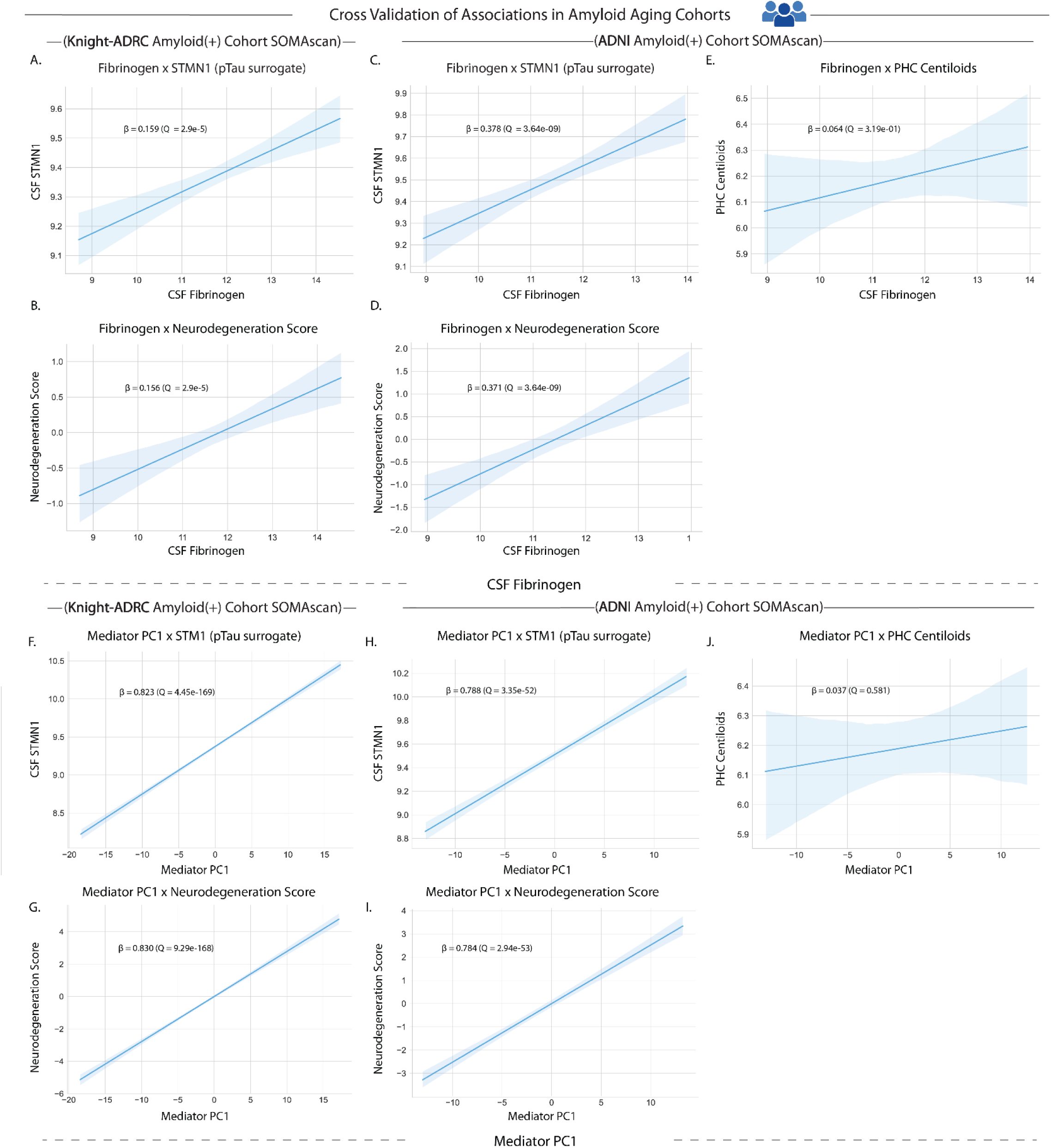
Thromboinflammatory markers maintain associations with tauopathy but not amyloidopathy in amyloid-positive disease states. **a-b,** Fibrinogen associations in Knight-ADRC amyloid-positive cohort (*N* = 752): STMN1 (pTau surrogate; β = 0.159, *Q* = 2.9e-5) and Neurodegeneration Score (β = 0.156, *Q* = 2.9e-5). **c-d,** Fibrinogen associations in ADNI amyloid-positive cohort (*N* = 265): STMN1 (β = 0.378, *Q* = 3.64e-9) and Neurodegeneration Score (β = 0.371, *Q* = 3.64e-9). **e,** No association between fibrinogen and PHC Centiloids in ADNI amyloid-positive cohort (β = 0.064, *Q* = 0.319). All CSF protein levels are log transformed. **f-g,** Mediator PC1 associations in Knight-ADRC amyloid-positive cohort (*N* = 752): STMN1 (β = 0.823, *Q* = 4.45e-169) and Neurodegeneration Score (β = 0.830, *Q* = 9.29e-168). **h-i,** Mediator PC1 associations in ADNI amyloid-positive cohort (*N* = 261-265): STMN1 (β = 0.788, *Q* = 3.35e-52) and Neurodegeneration Score (β = 0.784, *Q* = 2.94e-53). **j,** No association between Mediator PC1 and PHC Centiloids in ADNI amyloid-positive cohort (β = 0.037, *Q* = 0.581). All associations controlled for age and sex.

The mediator PC1 composite score similarly maintained strong associations with pTau markers and neurodegeneration in amyloid-positive states in both cohorts, while remaining independent of amyloid burden in ADNI (**Fig. 6f-j**; **Extended Data Fig. 7g-l**; **Supplementary Table 10**).

These consistent associations across amyloid-positive cohorts provide compelling evidence that thromboinflammation represents a distinct pathological axis that can drive tauopathy and neurodegeneration independently of amyloid pathology across the full disease spectrum.

## Discussion

In this study, we developed and applied an integrative analytical pipeline combining high-throughput CSF proteomics, mediation modeling, pathway enrichment, and multimodal phenotyping across three independent cohorts, totaling 1,655 participants, to uncover early signatures of brain vulnerability to immuno-vascular pathologies in aging (**Fig. 1**). We mapped a robust thromboinflammatory molecular network that mediates the effects of fibrinogen on neuronal synaptic vulnerability and tauopathy. Our findings suggest that coordinated thromboinflammation-associated changes may serve as early drivers of age-associated brain degeneration, potentially preceding amyloid deposition and clinical impairment. Importantly, this relationship remained significant through rigorous orthogonal methodological approaches and exhibited remarkable preservation across two independently collected external validation cohorts. Notably, the associations consistently maintained independence from biomarkers of amyloidopathy.

The implementation of a proteome-wide mediation framework in our discovery cohort identified seventy-two proteins that mediated the fibrinogen-synaptic degeneration relationship. Fifty-three protein mediators maintained their significance across all internal and external validation phases which included two biomarker platforms, namely targeted immunoassays and SOMAscan.

Enrichment analysis implicated pathways involved in neovascularization, vascular and immune regulation, and synaptic signaling networks. Positively-associated proteins showed robust enrichment for neovascularization/angiogenic activity, protein glycosylation, and NOTCH signaling pathways. Conversely, negatively associated mediators were enriched for vascular regulation, immune processes, and synaptic signaling networks. The presence of key genetic cSVD-associated proteins, including *NOTCH3* (mutated in CADASIL) and *FN1* (upregulated in both sporadic^38,39^ and genetic cSVD^36,37^), suggests that thromboinflammatory pathways may converge on mechanisms shared between sporadic aging and monogenic vascular diseases. This convergence is particularly intriguing given that CADASIL pathology involves aberrant NOTCH3 signaling and vascular smooth muscle cell dysfunction, which could contribute to the observed angiogenic dysregulation in our network^37^.

Prior mechanistic studies in rodents have demonstrated transcriptomic changes in fibrin-stimulated microglia that are captured by the mediator proteins reported in this work^11^. The concordance of prior mechanistic work in mice with the human-centered findings we report here provides strong evidence that thromboinflammatory networks are highly relevant to age-associated brain dysfunction and degeneration. These findings support a biological model whereby brain barrier dysfunction leads to the infiltration of blood proteins, activating brain-resident immune cells and promoting neuroinflammation-mediated synaptic degeneration. The identification of coordinated neovascularization responses suggests the formation of immature, hyperpermeable vessels that exacerbate barrier dysfunction, potentially through shared pathomechanisms with genetic cSVD, that create a self-amplifying pathophysiological cascade leading to widespread synaptic vulnerability.

Testing the clinical relevance of our findings, we identified significant associations between mediator proteins and compromised white matter microstructural integrity in long-range neuronal tracts, providing congruent evidence that links fibrinogen-centered thromboinflammatory signaling to structural brain changes that may precede clinically overt cognitive impairment. This evidence suggests that thromboinflammation may represent an early indicator of structural brain vulnerability, offering biomarker opportunities for identifying individuals at risk for adverse brain aging trajectories prior to clinical manifestation of neurodegenerative symptoms.

Importantly, fibrinogen consistently associated with biomarkers of tauopathy but not amyloidopathy across all cohorts investigated, suggesting a direct link between CNS thromboinflammation, brain barrier dysfunction, and tau-related neurodegeneration. This amyloid-independent association remained robust across the neurodegenerative continuum— from typical aging to established proteinopathies and persisted even in amyloid-positive individuals. These findings suggest that thromboinflammation represents a distinct pathophysiological risk axis that operates independently of the canonical amyloid cascade, directly contributing to tau-mediated neurodegeneration through immune and vascular mechanisms. Our study provides validation for vascular dysfunction as an early and amyloid-independent pathway for neurodegeneration in humans, with relevance to the fibrin-targeting antibody currently in clinical trials for AD^52^, and the numerous vascular therapeutics in development.

Several methodological limitations warrant acknowledgement. The cross-sectional design precludes temporal sequencing necessary for stronger causal inference, though we addressed this through robust statistical frameworks including bootstrapping and replication across independent datasets. Additionally, high-dimensional proteomics carries risk of false positives, which we minimized through stringent FDR correction and conservative interpretation restricted to proteins that consistently validated across both internal and external cohorts.

Overall, our study provides compelling evidence that subclinical changes to neurovascular unit integrity and toxic immune activation represent early pathologies driving brain dysfunction and degeneration. We identified a reproducible mediator network linking brain barrier permeability to structural brain outcomes, revealing thromboinflammation as a distinct pathophysiological axis that operates across aging and neurodegenerative disease states—critically, independent of amyloidopathy. This demonstrates that vascular dysfunction is not merely a byproduct of aging but may represent an initiating event in adverse brain aging and degeneration with wide-ranging consequences for brain health. This convergence of unbiased human data with previously established biological mechanisms demonstrated in rodents, positions CSF fibrinogen level as a quantifiable biomarker of brain barrier dysfunction with mechanistically relevant biology that connects age-related shifts in thromboinflammation to neuroglial vulnerability of the aging brain. Future therapeutics aimed at restoring the neurovascular interface^53^ may benefit from our findings, which identify molecular signatures and potential biomarkers that could be used for patient stratification and early assessment of therapeutic response.

Fibrinogen, and many of the identified mediator proteins represent druggable molecular targets, supporting future therapeutic investigations aimed at preserving cognitive function in vulnerable aging populations. These proteins may also offer complementary treatment targets to existing anti-amyloid therapeutics in amyloid-positive states. This comprehensive exploration of the CSF proteome has generated a compelling framework connecting cerebral small vessel disease and CNS immune activation with synaptic degeneration, tau hyperphosphorylation, and white matter alterations that may occur earlier than amyloid clearance abnormalities in the aging brain.

## Methods

### Study participants

#### UCSF Hillblom Longitudinal Aging Network (discovery cohort)

The discovery cohort comprised cognitively normal older adults enrolled in the observational Hillblom Longitudinal Aging Network and UCSF Alzheimer’s Disease Research Center studies conducted at the UCSF Memory and Aging Center. All study protocols were approved by the Institutional Review Board of the University of California, San Francisco, and written informed consent was obtained from all participants in accordance with the Declaration of Helsinki.

Participants underwent comprehensive cognitive and neuropsychiatric assessments administered by board-certified neurologists and neuropsychologists using standardized protocols. Cognitive normality was determined through clinical consensus conference review, with all participants maintaining functional independence (Clinical Dementia Rating Scale = 0) and demonstrating absence of amyloid pathology based on positron emission tomography imaging (18F-AV-45 standardized uptake value ratio ≤ 1.32^18^).

Cerebrospinal fluid samples underwent high-throughput proteomic analysis using the SomaLogic SomaScan v.4.1 platform (7,596 proteins) processed in a single analytical batch to minimize technical variance. A total of 66 cerebrospinal fluid samples from 66 participants constituted the discovery cohort for systematic biomarker identification and mediation analysis.

#### Knight Alzheimer’s Disease Research Center (validation cohort)

The Knight-ADRC represents a National Institute on Aging-funded longitudinal observational study conducted at Washington University School of Medicine in St. Louis under Institutional Review Board approval with written informed consent obtained from all participants. Research protocols conformed to established guidelines for human subjects’ protection and biospecimen handling.

For amyloid classification, participants were stratified based on cerebrospinal fluid Aβ_42_/Aβ_40_ ratios using established diagnostic thresholds previously validated within the Knight-ADRC cohort. Amyloid-negative status was defined by Aβ_42_/Aβ_40_ ratios above the established cutoff, while amyloid-positive classification required ratios below this threshold, indicating the presence of amyloid pathology independent of clinical symptom severity.

Proteomic analysis employed the SomaLogic SomaScan v.4.1 platform (7k) with standard normalization, calibration, and quality control procedures yielding relative fluorescence unit measurements. Cerebrospinal fluid samples from Knight-ADRC, ADNI, and other cohorts were processed using identical SomaScan protocols in the same analytical batch to ensure cross-cohort consistency and minimize technical variability.

Proteomic data from 482 amyloid-negative participants constituted the primary validation cohort, while 752 amyloid-positive participants comprised the exploratory cohort for pathway preservation analysis during established proteinopathy.

#### Alzheimer’s Disease Neuroimaging Initiative (validation cohort)

ADNI represents a longitudinal multicenter study designed for early Alzheimer’s disease biomarker development, with all data accessed through the ADNI database (https://adni.loni.usc.edu/) under established data use agreements and ethical approvals.

Cognitive assessment employed standardized diagnostic criteria with ADNI investigators systematically evaluating participants using the Mini-Mental State Examination (MMSE ≥ 24) and Clinical Dementia Rating Sum-of-Boxes Score (CDR-SB = 0 for cognitively normal classification). Additional inclusion criteria encompassed absence of clinical depression and preservation of activities of daily living to ensure cognitive normality determination.

Amyloid pathology status was determined through positron emission tomography imaging with PHC Centiloid quantification serving as the primary classification metric. Amyloid-negative participants were defined by PHC Centiloid values < 25, while amyloid-positive classification required PHC Centiloid values ≥ 25. This standardized threshold enabled precise amyloid burden quantification and direct assessment of fibrinogen independence from amyloid pathology.

Cerebrospinal fluid samples underwent proteomic analysis using the SomaLogic SomaScan v.4.1 platform (7k) with identical analytical protocols as employed for Knight-ADRC samples, ensuring cross-cohort analytical consistency. All samples were processed in the same analytical batch to minimize inter-cohort technical variance.

Proteomic data from 90 amyloid-negative, cognitively normal participants constituted the validation cohort, while 265 amyloid-positive participants comprised the exploratory cohort for amyloid independence assessment during established neurodegeneration.

#### Study design rationale

The multi-cohort analytical strategy was designed to provide systematic validation while minimizing potential sources of bias through independent replication across distinct research populations. Discovery analyses in the UCSF cohort established initial fibrinogen-synaptic marker associations, dual-cohort validation in Knight-ADRC and ADNI amyloid-negative populations confirmed reproducibility across independent samples, and exploratory analyses in corresponding amyloid-positive cohorts assessed pathway preservation and amyloid independence during established neurodegeneration. Specifically, we developed a novel five-stage analytical framework spanning the full spectrum from normal aging to amyloid-positive AD (**Fig.1**). (1) establishing fibrinogen-synaptic marker relationships in amyloid-negative, cognitively typical older adults (UCSF Hillblom, *N* = 66; Knight ADRC, *N* = 482; ADNI, *N* = 73); (2) identifying molecular mediators through high-dimensional proteomic mediation analysis across discovery and validation cohorts; (3) determining cellular sources via single-cell transcriptomic mapping; (4) construct validation by testing the association of mediators to clinical outcomes (UCSF Hillblom cohort); and (5) testing the relevance of captured biology to amyloid-positive brain aging states (Knight ADRC, *N* = 752; ADNI, *N* = 472). This comprehensive approach ensured robust characterization of thromboinflammatory mechanisms across the complete spectrum of brain aging and neurodegeneration while maintaining analytical rigor through standardized proteomic platforms and consistent biomarker classification criteria.

### CSF biomarker measurements

#### Targeted immunoassays

CSF samples and assays were collected and completed by an independent group as described in previous publications^54,55^. Briefly, lumbar punctures were conducted following a 12-hour fast. CSF was processed in accordance with standard protocols. The levels of Aβ_42_, Aβ_40_, and pTau-181 were measured using the Lumipulse platform. The quantification of synaptic proteins (NG36, GAP43, SNAP25) has also been previously detailed^35,36^. CSF fibrinogen was assayed in-house via enzyme-linked immunosorbent assay (ELISA) according to manufacturer’s standard protocol (Abcam ab108841; 1:200 dilution). All biomarker measurements were log2- transformed prior to analysis to normalize distributions and stabilize variance.

Missing biomarker values (2.8% fibrinogen, 22.5% SNAP25, < 2% for other quantified proteins) were determined to be missing completely at random (MCAR) using Little’s test and underwent iterative imputation using the IterativeImputer algorithm from scikit-learn^56,57^. The imputation model incorporated complete proteomic profiles and available protein markers as predictors, employed 5 iterations with different random states to generate multiple imputed datasets, and applied bootstrapping with posterior sampling to capture uncertainty in imputed values.

#### Unbiased high-dimensional proteomics

The SomaLogic SomaScan assay (https://somalogic.com/), which uses slow off-rate modified DNA aptamers (SOMAmers) to bind target proteins with high specificity, was used to quantify relative concentrations of human proteins in CSF. The v.4.1 (∼7,000 proteins) assay was used for all cohorts. Standard SomaLogic normalization, calibration and quality control were performed on all samples, resulting in protein measurements in relative fluorescence units. The resulting values were log₁₀ normalized as the assay had an expected log-normal distribution. No cohort batch corrections were applied.

Quality control procedures included assessment of coefficient of variation across technical replicates, with 95% of proteins demonstrating < 20% CV. Proteomics data underwent systematic filtering to remove failed samples (RowCheck = “FLAG”) and low-quality protein measurements (ColCheck = “FLAG”), resulting in 7,168 high-quality protein measurements for downstream discovery analysis.

#### Cross-platform validation and surrogate marker selection

To validate findings across independent cohorts with different measurement platforms, systematic cross-platform comparisons were performed between specific proteins measured via immunoassays and SOMAscan measurements in the discovery cohort. Strong concordance was observed for fibrinogen, GAP43, and NRGN (*Q* < 5.00e-12). For proteins with weaker cross-platform agreement (pTau and SNAP25), functionally related surrogate markers were identified based on biological relevance and correlation strength.

STMN1 (stathmin-1), a microtubule-regulating phosphoprotein, served as a biological surrogate for pTau based on robust correlation with pTau (*P* = 2.68e-16) and preserved association with fibrinogen across discovery and validation cohorts. SHANK3 (SH3 and multiple ankyrin repeat domains 3), a key scaffolding protein in synaptic signaling, functioned as a surrogate for SNAP25 with strong correlation to SNAP25 (*P* = 6.01e-15) and maintained fibrinogen relationships across cohorts.

Surrogate marker specificity was assessed through sensitivity analysis using 1,000 bootstrap iterations with randomly selected proteins matched on expression level and abundance parameters (**Extended Data Figure 8**). The observed associations were not recapitulated by random proteins, confirming functional relevance and biological specificity of chosen surrogates.

### Statistical analyses

#### Software and data processing

All statistical analyses were conducted using Python (v3.10.11) leveraging pandas^58^ and NumPy^59^ for data processing, matplotlib^60^ and seaborn^61^ for visualization, and statsmodels^62^ for statistical modeling. Additional analyses employed scikit-learn^57^ for machine learning approaches.

#### Weighted protein co-expression network analysis (WPCNA)

WPCNA was performed using the PyWGCNA package^20,63^ to identify clusters of co-expressed proteins within the CSF proteome. Soft-thresholding powers (β = 15-26) were evaluated using scale-free topology criteria to optimize network construction. A signed network with minimum module size of 5 proteins was constructed to identify biologically coherent protein modules.

Module membership (correlation with module eigengene) and cross-cohort preservation statistics were calculated to assess network stability across independent cohorts.

#### Linear regression analysis

The ordinary least squares (OLS) function from statsmodels^62^ was used to assess linear associations between protein levels and synaptic markers. All continuous variables underwent log transformation prior to analysis to normalize distributions and meet linear model assumptions. Models were systematically adjusted for age and sex as primary covariates, with additional covariates included as specified in individual analyses.

For proteome-wide association testing, standardized beta coefficients were calculated to enable cross-protein comparison of effect sizes. Multiple hypothesis testing correction was applied using the Benjamini-Hochberg method with significance threshold set at 5% false discovery rate (*Q* < 0.05). Effect sizes were interpreted as standardized coefficients representing the standard deviation change in outcome per standard deviation change in predictor.

#### Mediation analysis

To identify molecular mediators linking fibrinogen to synaptic markers, mediation analysis was conducted using the Baron and Kenny framework^32^. The mediation analysis included three critical pathways: (1) assessment of the total effect of fibrinogen on synaptic markers (c-path); (2) evaluation of fibrinogen’s effect on potential mediators (a-path); and (3) estimation of the mediator’s effect on synaptic markers adjusted for fibrinogen (b-path).

Indirect effects were quantified using the product-of-coefficients method (a x b), with statistical significance determined through nonparametric bootstrapping (1,000 iterations). Bootstrap confidence intervals were calculated using bias-corrected and accelerated (BCa) methods to provide robust inference for indirect effects. E-values were calculated to quantify the strength of unmeasured confounding required to explain away observed mediation effects.

Mediation screening was conducted across all 7,168 QC-passed proteins in the discovery cohort, with validation performed in independent cohorts using identical analytical frameworks. Only proteins demonstrating consistent mediation across all cohorts and outcome measures were retained for downstream biological characterization.

#### Principal component analysis

Principal component analysis was performed using the PCA function from scikit-learn with data preprocessing using StandardScaler to normalize protein expression levels. PCA was applied to: (1) generate composite neurodegeneration scores from correlated synaptic biomarkers (pTau, GAP43, NRGN, SNAP25); and (2) explore joint associations of validated mediator proteins with neuroimaging outcomes.

The first principal component (PC1) was extracted as a summary measure, with loadings interpreted to understand the contribution of individual proteins to the composite score. Variance explained by each component was calculated, with PC1 typically explaining > 85% of total variance in synaptic biomarker combinations.

#### Cross-cohort validation and amyloid independence testing

Systematic cross-cohort validation was performed to assess generalizability of thromboinflammation-synaptic relationships across independent populations. Linear regression models with identical covariate structures were applied across all cohorts, with effect size consistency evaluated using standardized beta coefficients and confidence interval overlap.

Amyloid independence was assessed through multiple approaches: (1) restriction of primary analyses to amyloid-negative participants (18F-AV-45 SUVR ≤ 1.32^18^ or PHC Centiloids < 25^19^); (2) direct testing of fibrinogen associations with amyloid biomarkers (Aβ_42/40_ ratio, PHC Centiloids); and (3) preservation of associations in amyloid-positive cohorts to assess pathway persistence across the neurodegeneration spectrum.

#### Pathway enrichment analysis

Gene Ontology enrichment analyses were performed using EnrichR across eight curated databases: GO Biological Process (2025), GO Cellular Component (2025), GO Molecular Function (2025), Jensen Disease (Curated and Experimental; 2025), Reactome (2024), SynGO (2024), and WikiPathways (2024). The SomaScan 7k protein panel served as the background gene set for enrichment calculations.

Enrichment analyses were conducted separately for proteins positively and negatively associated with neurodegeneration to identify distinct pathway signatures. Statistical significance was assessed using un-adjusted *P* < 0.05.

#### Single-cell transcriptomic mapping

To investigate cellular sources of validated mediator proteins, single-cell RNA sequencing data from *ex vivo* human brain tissue was analyzed using one-versus-rest differential expression analysis^29^. Cell type-specific expression patterns were evaluated across principal neurovascular and glial cell types, including endothelial cells, pericytes, smooth muscle cells, oligodendrocytes, microglia/macrophages, and fibroblasts.

Expression enrichment was quantified using log_2_ fold change values and the percentage of cells expressing each gene within specific cell types. Proteins were classified as cell type-enriched based on preferential expression patterns and functional relevance to neurovascular unit biology.

#### Reporting and transparency

All analyses followed STROBE guidelines for observational studies64. Multiple testing corrections were applied consistently across all analyses, with both raw and adjusted p-values reported. Effect sizes with confidence intervals were prioritized over significance testing alone. Missing data patterns were systematically evaluated and addressed through appropriate imputation methods where justified.

Sensitivity analyses were conducted to assess robustness of findings to analytical choices, including alternative imputation strategies, covariate selection, and outlier treatment. All code and analysis pipelines are made available for reproducibility at https://github.com/mssm-elahi-lab/thromboinflammation.

#### Reporting summary

Further information on research design is available in the Nature Portfolio Reporting Summary linked to this article.

## Data availability

All data are available upon reasonable request with formal applications submitted to respective cohort committees to protect patient-sensitive data. UCSF data can be requested from the UCSF Hillblom Longitudinal Aging Network at https://memory.ucsf.edu/research-trials/professional/open-science. Knight-ADRC proteomics data are available upon request at https://knightadrc.wustl.edu/. ADNI data can be requested at https://adni.loni.usc.edu/. EnrichR can be accessed at https://maayanlab.cloud/Enrichr/. Single-cell RNA sequencing data from Wälchli et al. are publicly available at https://cells.ucsc.edu/brain-vasc/unsorted/adult-control/adult-control-brain-unsorted.rds.

## Code availability

All code for data processing and analyses is available at https://github.com/mssm-elahi-lab/thromboinflammation/. The CSF biomarker data processing pipeline and statistical analyses are implemented in Python and R with detailed documentation for reproducibility.

## Acknowledgements

This research was supported by the Edward and Pearl Fein gift to F.M.E and K.A.; the National Institute on Aging and Department of Veterans Affairs IK2CX002180, Larry L. Hillblom Foundation 2019A012SUP and New Vision Research to F.M.E; the Dolby Family Fund, the Simon Family Trust, Brightfocus, NIH/NIA RF1 AG064926, and NIH/NINDS R35 NS097976 to K.A; NIH R01NS122888, UH3NS106899, U24NS122732, US VA 1I01RX002245, I01RX002787, I01BX005871, I50BX005878, Craig H. Neilsen Foundation, and Wings for Life Foundation to A.R.F. The validation of results included data shared by the Alzheimer’s Disease Neuroimaging Initiative (ADNI) (National Institutes of Health Grant U19 AG024904).

## Author contributions

FME conceived the study. HLR, JNK, ARF, and FME designed the detailed analytical plan. NB, AMK, KA and FME generated fibrinogen levels from CSF. JNK and HLR performed the biostatistical analyses and generated figures. JNK, HLR, KA, ARF, and FME critically discussed and interpreted the results. KC, BLM, CC, and JK shared existing data and JK shared samples. HLR, JNK, ARF, and FME drafted the manuscript. All authors critically read and provided edits. FME and ARF provided funding.

